# Phase-separated TRB-PRC2 aggregates contribute to Polycomb silencing in plants

**DOI:** 10.1101/2022.03.27.485997

**Authors:** Hua Xuan, Yue Liu, Jingze zhao, Nan Shi, Yanzhuo Li, Yulu Zhou, Limin Pi, Shaoqing Li, Guoyong Xu, Hongchun Yang

## Abstract

Local accumulation of Polycomb Repressive Complex 2 (PRC2) is essential to gene silencing. Liquid-liquid phase separation (LLPS) mechanism is emerging as a paradigm to concentrate transcriptional machinery for effective gene regulation. Here, we elucidate that a rice single Myb transcription factor TRBF2 forms phase-separated droplets, which aggregate with PRC2 through direct protein interaction. Furthermore, TRB1, the closest homolog of TRBF2 in *Arabidopsis*, also forms phase-separated aggregates with PRC2. Mutants of *TRBF2* and PRC2 component *CLF* display similar developmental defects, share common differentially expressed genes, and reduced H3K27me3 chromatin regions. Chromatin immunoprecipitation analysis supports that TRBF2 concentrates PRC2 at target loci to promote H3K27me3 deposition. Therefore, we propose that the aggregation of the plant-specific TRBs with PRC2 by the LLPS mechanism contributes to Polycomb silencing.

**One-Sentence Summary:** The phase-separated plant-specific single Myb transcription factor aggregates with PRC2 to facilitate Polycomb silencing.

## Main Text

The evolutionarily conserved Polycomb Repressive Complex 2 (PRC2) triggers and maintains gene silencing by catalyzing histone 3 lysine 27 tri-methylation (H3K27me3), hence, controlling development and differentiation in higher eukaryotes. Studies have revealed that the recruiting and accumulation of PRC2 at target loci are essential for Polycomb silencing and epigenetic inheritance (*1–7*). Several transcription factors have been identified, which guide PRC2 to target loci through direct protein interaction in *Drosophila*, mammals, and dicot plant *Arabidopsis* (*4, 5, 8–13*). On the other hand, it has long been speculated that the formation of the Polycomb body is critical for gene silencing (*14, 15*). Whether PRC2 forms nuclear bodies and the role of transcription factors on this process have remained an enigma.

PRC2 plays a crucial role in controlling monocot plant rice (*Oryza sativa*) development (*16–20*), but the regulatory role of transcription factors on PRC2 function is largely unknown. To this end, the core PRC2 component EMF2b (EMBRYONIC FLOWER 2b) was used as the bait to screen a library containing 1,588 rice transcription factors by yeast two-hybrid assay (Y2H) (Fig. S1 and table S1). A plant-specific single Myb transcription factor TRBF2 (Telomere Repeat-binding Factor 2) was identified (Fig. 1A and fig. S1C). We further found TRBF2 also interact with the other Su(z)12 component EMF2a, and both E(z) components CLF (CURLY LEAF) and SET1 (SETDOMAIN 1), but not ESC components FIE1 (FERTILIZATION-INDEPENDENT ENDOSPERM 1) and FIE2 in yeast cells (Fig. S2A). To test whether TRBF2 associates with PRC2 in *planta*, tagged versions of CLF (CLF-HA) and EMF2b (EMF2b-HA) were co-expressed with TRBF2 (TRBF2-Venus) in *Nicotiana benthamiana* (*N. ben*) leaf. The nuclear proteins were extracted and subjected to co-immunoprecipitation. CLF-HA and EMF2b-HA were co-precipitated with TRBF2-Venus (Fig. S2, B and C). Further, we also detected the interaction between TRBF2 and CLF in rice (Fig. 1B). These data displayed that the rice transcription factor TRBF2 forms a protein complex with PRC2. TRBF2 homologs in *Arabidopsis* also associate with PRC2, thereby, recruiting PRC2 to target loci (*13, 21*). Together, these suggest that the association of plant-specific telomere repeat-binding factor with PRC2 is likely to be conserved across monocots and dicots.

**Fig. 1.**
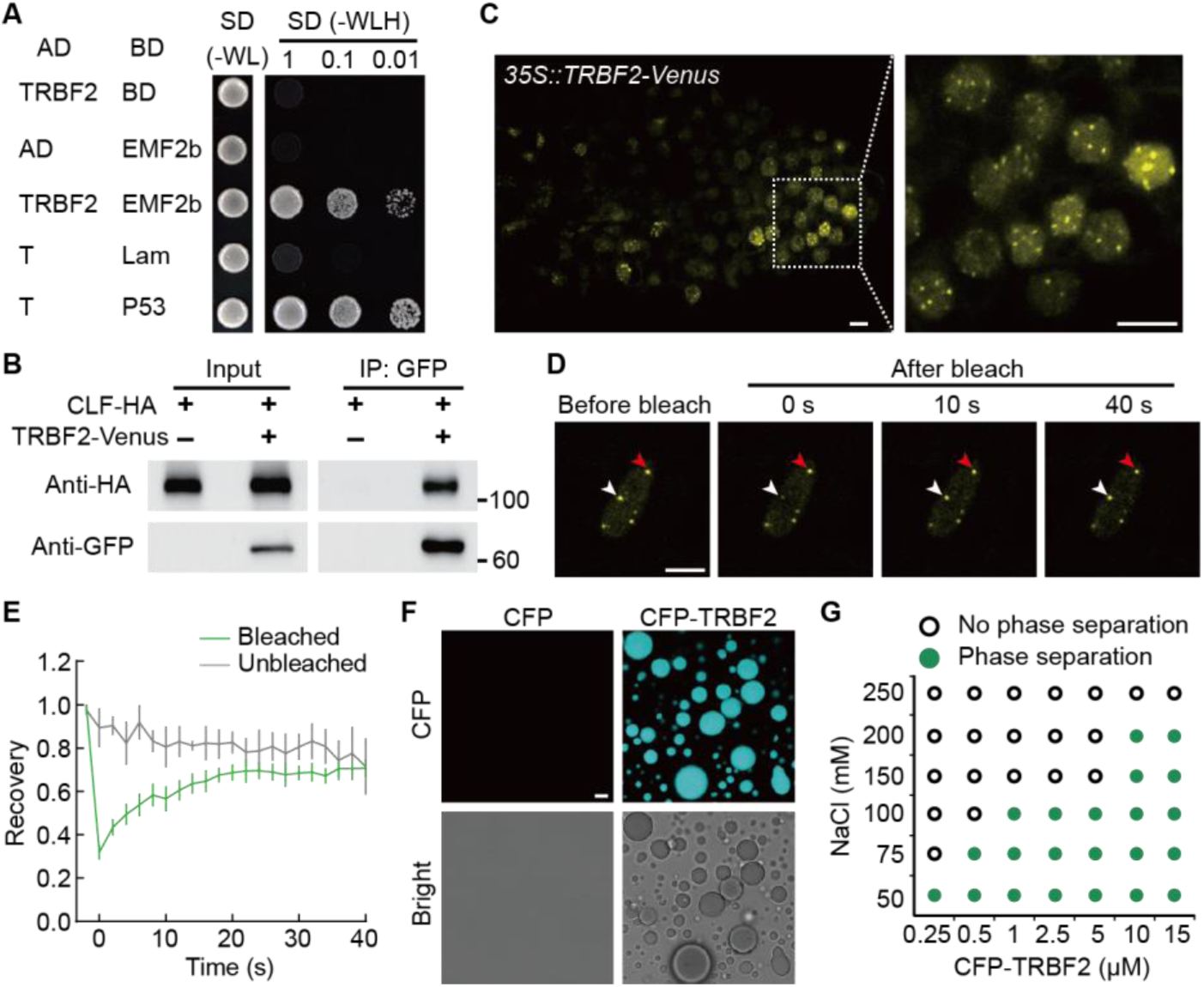
Rice TRBF2 associates with PRC2 and exhibits phase separation characteristics. **(A)** TRBF2 interacts with EMF2b in yeast cells. TRBF2 and EMF2b were fused with AD and GAL4-BD domain respectively. Paris of indicated plasmids were co-transformed and grown on indicated solid media. Representative pictures from two independent experiments are shown. (**B**) CLF-HA is co-immunoprecipitated with TRBF2-Venus in rice. Hemagglutinin (HA)-tagged CLF (CLF-HA) and Venus-tagged TRBF2 (TRBF2-Venus), or CLF-HA along served as a negative control, were expressed in rice protoplasts. Protein was extracted and subjected to co-immunoprecipitation. Samples before and after immunoprecipitation were detected by anti-HA and anti-GFP. Images are representative of two independent experiments. (**C**) Fluorescence microscopy of rice lateral root tip cells that express TRBF2-Venus. A stable transgenic line carrying *35S::TRBF2-Venus* was grown under hydroponic culture conditions. The 12-day old seedling was used in the analysis. Scale bars, 5 μm. Images are representative of three independent experiments. (**D**) FRAP of TRBF2-Venus in rice lateral root tip cells carrying the *35S::TRBF2-Venus* transgene. Time 0 indicates the time of the photobleaching pulse. The white arrow indicates the nuclear body that is bleached. The red arrow indicates the unbleached nuclear body used as the control. Pictures are representative of eight independent experiments. Scale bar, 5 μm. (**E**) FRAP recovery curves of TRBF2-Venus (green line). The gray line displays fluorescence variation of unbleached nuclear bodies. Values are mean ± s.d. (n = 8). (**F**) Images showing that CFP-TRBF2 forms phase-separated droplets at the concentration of 5 μM. NaCl concentration is 50 mM. Images are representative of three independent experiments. Scale bar, 5 μm. (**G**) Diagram showing the phase formation of CFP-TRBF2 at different concentrations of CFP-TRBF2 and NaCl. Images are representative of three independent experiments.

To evaluate the subcellular localization of TRBF2, stable transgenic rice carrying the fluorescence protein-tagged TRBF2 (TRBF2-Venus) was generated. The TRBF2-Venus fluorescence was mainly localized in the nucleus with several intense foci in the lateral root tip cells (Fig. 1C). TRBF2-Venus was also transiently expressed in rice protoplasts and *N. ben* leaf epidermal cells. Similar localization signals were also observed (Fig. S3, A and B). The morphology of these foci looked similar to that of condensates formed by transcriptional regulators reported previously (*22–24*). Interestingly, we further found these TRBF2-Venus foci have liquid-like properties by fluorescence recovery after photobleaching (FRAP) assay. After photobleaching, the fluorescence of TRBF2-Venus foci gradually recovered within 1 minute both in rice lateral root tip cells and *N. ben* leaf epidermal cells (Fig. 1, D and E, fig. S3, C and D, and movies S1 and S2), indicating a fast protein exchange between TRBF2-Venus condensates and surrounding nucleoplasm. Taken together, these observations displayed that TRBF2 exhibits the characteristics of biomolecular condensates driven by liquid-liquid phase separation (LLPS) in the nucleus, and behaves similarly in stable transgenic rice and transient expression systems.

We next examined whether TRBF2 can be phase-separated individually, recombinant CFP-TRBF2 was expressed in *E. coli* BL21 cells and further purified by affinity chromatography. The protein was eluted by elution buffer with 500 mM NaCl (Fig. S4, A and B). Diluting the NaCl concentration to 50 mM turned the protein solution turbid, and the turbidity became more manifest with increasing CFP-TRBF2 concentration, whereas the solution with CFP alone remained clear (Fig. S4C). Checking the solution under the microscope, we observed spherical droplets of CFP-TRBF2 (Fig. 1F). The formation of CFP-TRBF2 droplets could be observed at a very low concentration (0.25 μM) (Fig. S4D). The number and size of the droplets increased with increasing CFP-TRBF2 concentration, declined with the increase of NaCl concentration (Fig. 1G and fig. S4D). The CFP-TRBF2 signal within droplets recovered after photobleaching, indicating CFP-TRBF2 molecules diffused rapidly within droplets (Fig. S4, E and F, and movie S3). Approaching droplets could fuse into big puncta upon contact (Fig. S4G and movie S4). These data collectively support that TRBF2 exhibits the characteristics that are consistent with phase separation *in vivo* and undergoes LLPS *in vitro*.

Phase-separated protein condensates can attract its interacting protein to facilitate gene regulation (*23, 25*). For example, the Pc component of *Drosophila* PRC1 could be enriched by the Mini-Ph (amino acid 1,289–1,577 of Polyhomeotic) – chromatin condensates to stimulate Polycomb silencing (*25*). As TRBF2 forms a protein complex with PRC2, we next asked whether TRBF2 condensates affect PRC2 behavior. CLF was expressed alone or co-expressed with TRBF2 in rice protoplasts. Individually expressed CLF-YFP displayed even distribution in nuclei (Fig. S5A). However, several CLF-YFP nuclear bodies were observed once co-expressed with TRBF2-CFP (Fig. S5A). And these nuclear bodies were co-localized with TRBF2-CFP condensates (Fig. S5, A and B). A similar characteristic was also observed in *N. ben* leaf epidermal cells when CLF-YFP was co-expressed with TRBF2-CFP (Fig. 2, A and B). EMF2b-YFP was also co-expressed with TRBF2-CFP in *N. ben* leaf. Several EMF2b-YFP nuclear bodies were observed and co-localized with TRBF2-CFP condensates (Fig. S5, C and D). We only observed the PRC2 condensates when PRC2 components were co-expressed with TRBF2-CFP *in planta*, but not individually expressed (Fig. 2, A and B, and fig. S5, A to D), indicating PRC2 condensation is TRBF2 dependent. Next, we would like to address whether PRC2 could form biomolecular condensates *in vivo*. Transgenic rice carrying *35S::CLF-Venus* was generated and uneven CLF-Venus fluorescence was observed. We observed several relatively brighter dots in the nucleus of lateral root tip cells (Fig. 2C), indicating PRC2 forms condensates *in vivo*. However, we could not study the dynamics of CLF-Venus puncta *in vivo* because of the limitation of low expression. So we employed the transient expression system to address this question. CLF-YFP and TRBF2-CFP were co-expressed in *N. ben* leaf epidermal cells, and CLF-YFP nuclear bodies were photobleached by a 514 nm laser. A certain recovery of CLF-YFP fluorescence was observed, and the unbleached TRBF2-CFP at the same region remained stable (Fig. 2, D and E, and movie S5), indicating that CLF-YFP could be redistributed between puncta and surrounding nucleoplasm. Together, these observations suggest that PRC2 can be attracted and concentrated by TRBF2 condensates to form nuclear bodies, which display characteristics driven by LLPS in *planta*.

**Fig. 2.**
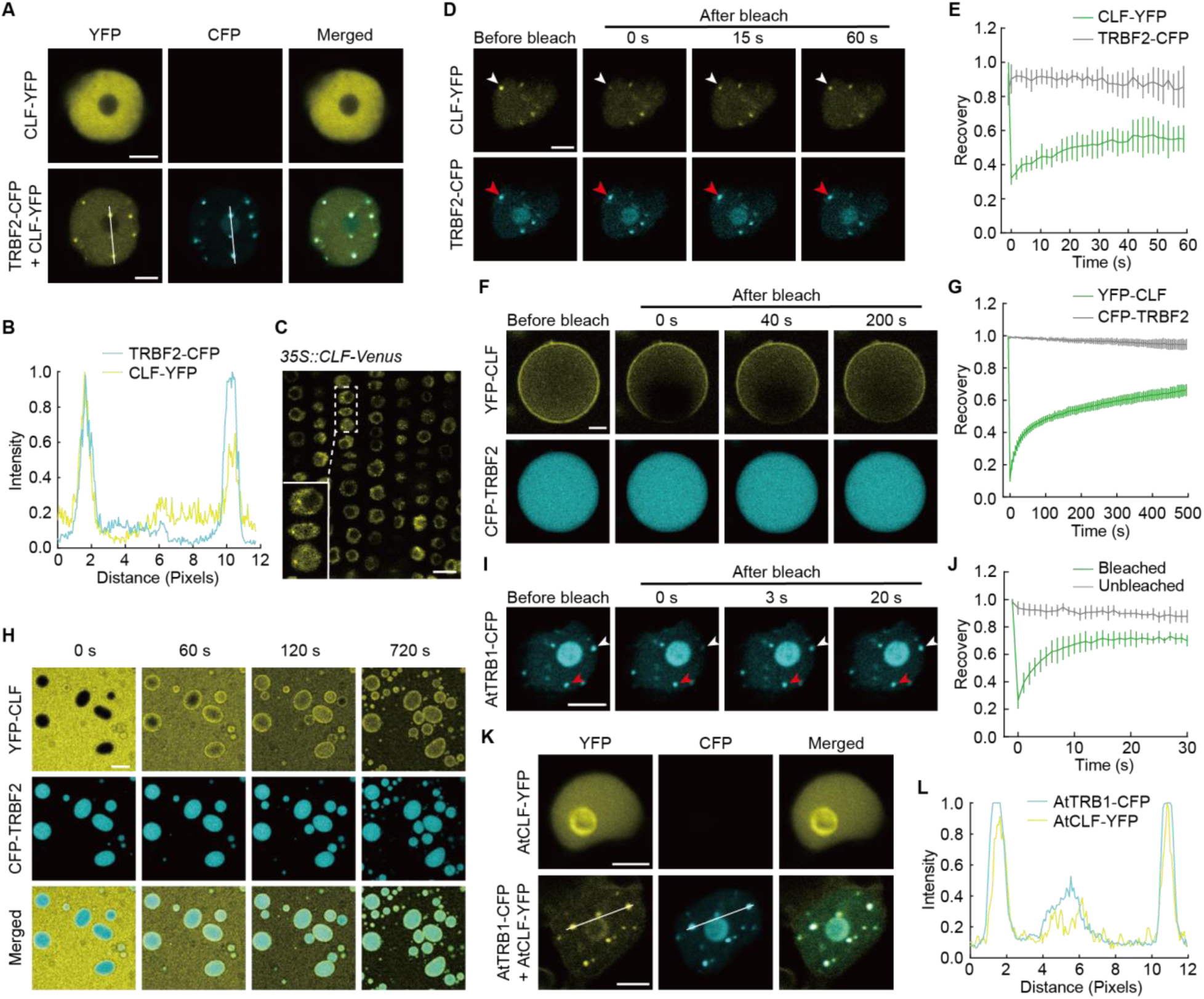
Plant TRB condensates attract PRC2 to form phase-separated aggregates *in vivo* and *in vitro*. (**A**) Images showing CLF-YFP forms nuclei bodies colocalized with TRBF2-CFP condensates when co-expressed with TRBF2-CFP in *N. ben* leaf epidermal cells. Top panel, express CLF-YFP alone; bottom panel, co-express CLF-YFP with TRBF2-CFP. Scale bars, 5 μm. Images are representative of three independent experiments. (**B**) Analysis of colocalization of TRBF2-CFP (cyan line) and CLF-YFP (yellow line). The fluorescence intensities (y-axis) are plotted along the while line in (A) (x-axis). Images are representative of three independent experiments. (**C**) Fluorescence microscopy of rice lateral root tip nuclei that express CLF-Venus. Inset represents a zoomed in view of the dash-lined box. A stable transgenic line carrying *35S::CLF-Venus* was grown under hydroponic culture conditions. The 6-day old seedling was used in the analysis. Scale bars, 10 μm. Images are representative of three independent experiments. (**D**) FRAP of CLF-YFP when co-expressed with TRBF2-CFP in *N. ben* leaf epidermal cells. Top panel, YFP channel; bottom panel, CFP channel. Time 0 indicates the time of the photobleaching pulse. The white arrow indicates the nuclear body that is bleached. The red arrow indicates the unbleached TRBF2-CFP of the same nuclear body. Pictures are representative of eight independent experiments. Scale bar, 5 μm. (**E**) FRAP recovery curves of CLF-YFP (green line) when co-expressed with TRBF2-CFP in *N. ben* leaf epidermal cells. The gray line displays the fluorescence of unbleached TRBF2-CFP nuclear bodies. Values are mean ± s.d. (n = 8). (**F**) FRAP of YFP-CLF droplets when mixed with CFP-TRBF2 *in vitro*. Top panel, YFP channel; bottom panel, CFP channel. Time 0 indicates the time of the photobleaching pulse. CFP-TRBF2 protein 2.5 µM, YFP-CLF protein 0.25 µM and NaCl 50 mM. Images are representative of ten independent experiments. Scale bar, 2 μm. (**G**) FRAP recovery curves of YFP-CLF (green line) when mixed with CFP-TRBF2 *in vitro*. The gray line displays fluorescence variation of CFP-TRBF2 at the same region. Values are mean ± s.d. (n = 10). (**H**) Images showing progression of YFP-CLF droplets formation when mixed with CFP-TRBF2 *in vitro*. Time 0 indicates the time of the mixing. CFP-TRBF2 protein 2.5 µM, YFP-CLF protein 0.25 µM, and NaCl 50 mM. Images are representative of three independent experiments. Scale bar, 20 µm. (**I**) FRAP of AtTRB1-CFP in *N. ben* leaf epidermal cells. Time 0 indicates the time of the photobleaching pulse. The white arrow indicates the nuclear body that is bleached. The red arrow indicates the unbleached nuclear body used as the control. Pictures are representative of seven independent experiments. Scale bar, 5 μm. (**J**) FRAP recovery curves of AtTRB1-CFP (green line) in *N. ben* leaf epidermal cells. The gray line displays fluorescence variation of unbleached nuclear bodies. Values are mean ± s.d. (n = 7). (**K**) Images showing AtCLF-YFP forms nuclei bodies colocalized with AtTRB1-CFP condensates when co-expressed with AtTRB1-CFP in *N. ben* leaf epidermal cells. Top panel, express AtCLF-YFP alone; bottom panel, co-express AtCLF-YFP with AtTRB1-CFP. Scale bars, 5 μm. Images are representative of three independent experiments. (**L**) Analysis of colocalization of AtTRB1-CFP (cyan line) and AtCLF-YFP (yellow line). The fluorescence intensities (y-axis) are plotted along the while line in (K) (x-axis). Images are representative of three independent experiments.

We further tested TRBF2-PRC2 aggregates formation *in vitro*. A small amount of low concentration YFP-CLF was purified (Fig. S6, A and B). YFP-CLF could not form condensate alone (Fig. S6C). CFP-TRBF2 could not induce solo YFP to form condensate either (Fig. S6C). However, YFP-CLF puncta were observed when mixed with CFP-TRBF2, and co-localized with CFP-TRBF2 condensates (Fig. S6C). Further, part of YFP-CLF puncta was bleached, gradual recovery of YFP-CLF was detected, whereas unbleached CFP-TRBF2 at the same region was unchanged (Fig. 2, F and G, and movie S6). To study the progression of YFP-CLF puncta formation, YFP-CLF and CFP-TRBF2 were mixed under the confocal microscope, images were taken immediately. YFP-CLF could be attracted and concentrated by CFP-TRBF2 condensates and reduced YFP-CLF signal outside of CFP-TRBF2 droplets was also observed upon time progression (Fig. 2H and movie S7). The sedimentation assay displayed that the YFP-CLF was enriched in the condensates together with TRBF2 (Fig. S6, D and E). YFP-EMF2b also can be attracted by TRBF2 condensates to form LLPS puncta and enriched within the condensates *in vitro* (Fig. S6, C and F). Plant-specific telomere repeat-binding factors are widely distributed in monocots and dicots (*21*). *Arabidopsis* TRBs were proven to be associated with PRC2 (*13*). So we tested whether the *Arabidopsis* TRBs have similar characteristics as rice TRBF2. AtTRB1, the closest homolog of TRBF2 in *Arabidopsis*, was used in the following assays (*21*). First, AtTRB1 was expressed in *N. ben* leaves. Several AtTRB1 foci were observed (Fig. 2I), which is consistent with the previous report (*26*). We performed a FRAP assay and found that after photobleaching, AtTRB1 redistributed rapidly from the unbleached area to the bleached area (Fig. 2, I and J, and movie S8). And the purified CFP-AtTRB1 could form spherical droplets *in vitro* (Fig. S6C). These data displayed that the AtTRB1 behaves like LLPS condensates in *planta* and *in vitro* as rice TRBF2. Next, we examined whether AtTRB1 was able to attract the *Arabidopsis* PRC2 component to its condensates. AtCLF-YFP was co-expressed with AtTRB1-CFP in *N. ben* leaves. We found AtCLF-YFP was concentrated in AtTRB1-CFP condensates (Fig. 2, K and L). Consistently, AtCLF was also attracted by AtTRB1 droplets *in vitro* (Fig. S6, B and C). Together, our data support that PRC2 is attracted by protein condensates of transcription factor TRB and forms LLPS aggregates with TRB in rice and *Arabidopsis*.

To explore the function of TRBF2 in rice, three independent mutants were generated by CRISPR-Cas9 method (Fig. S7A). Different from the functional redundancy of *Arabidopsis* TRBs (*13*), all the single mutants of *TRBF2* displayed obvious developmental defects, such as severe dwarf because of reduced length of internodes, reduced panicle length, and decreased numbers of primary and secondary panicle branches (Fig. 3A and fig. S7), suggesting critical roles of TRBF2 in controlling rice development. We further investigated the molecular phenotype of *trbf2* by RNA-seq transcriptome profiling. A total of 1,463 differentially expressed genes (DEGs) including 782 up- and 681 down-DEGs were identified (Fig. 3B and table S2). Down-DEGs displayed relatively similar expression levels as unchanged genes in wild type (WT) (Fig. 3C). However, most of the up-DEGs were not or low expressed in WT (Fig. 3C), suggesting a repression role of TRBF2 on these genes. As TRBF2 forms LLPS aggregates with PRC2, we next probed H3K27me3 levels in *trbf2* by cut&tag (*27*). In total, 12,378 H3K27me3 enriched regions were identified in *trbf2* (Fig. 3D). More than half of these regions were located at the 5’ end of genes near the transcription start site (TSS) (Fig. S8A). We further quantified the level of each peak using the Diffbind program (*28*). Most H3K27me3 enriched regions were common between WT and *trbf2* (unchanged H3K27me3). Still, a lower mean level of all H3K27me3 regions was observed compared to WT (Fig. 3, D and E), indicating TRBF2 is required for H3K27me3 deposition at a subset of PRC2 targets. 2,286 lower (lost H3K27me3 in *trbf2*) and 560 higher regions (gained H3K27me3 in *trbf2*) respectively were detected (Fig. 3, D to F, and table S3). Most of the changed H3K27me3 peaks were overlapped with genes (Fig. S8B). The mean level of H3K27me3 at genes in *trbf2* was also lower than in WT, and 2,083 and 422 genes significantly lost or gained H3K27me3 in *trbf2* (Fig. S8, C and D, and table S3). A significant amount of altered H3K27me3 genes were overlapped with DEGs in *trbf2* (Fig. 3G). Thus, TRBF2 is required for H3K27me3 distribution and gene regulation at a subset of the PRC2 targets.

**Fig. 3.**
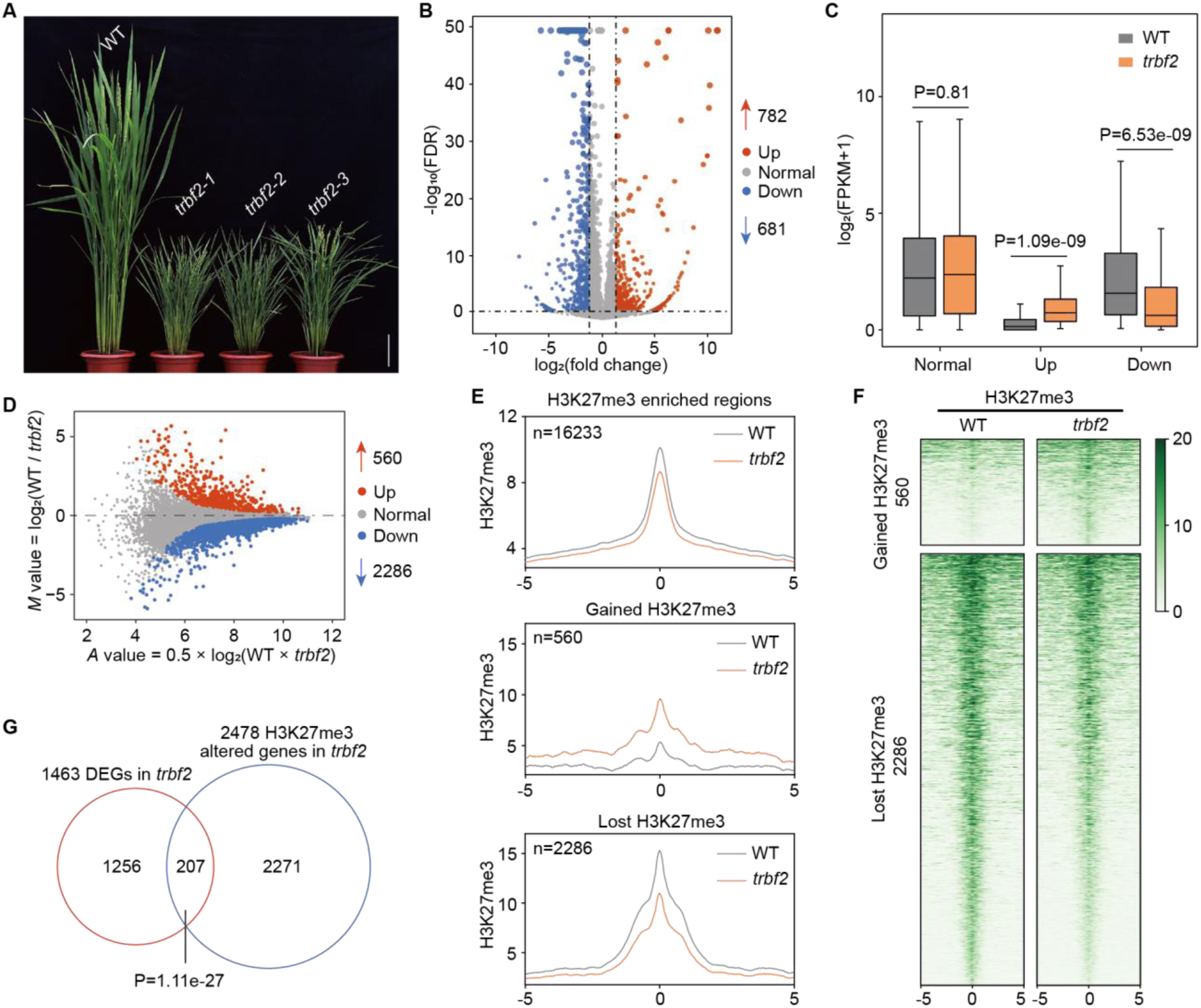
Analysis of *TRBF2* mutants defective in gene expression and H3K27me3 deposition at the genome-wide level. (**A**) Representative plants of *trbf2* compared to WT at the heading stage. Scale bar, 10 cm. (**B**) Volcano plot showing differentially expressed genes between WT and *trbf2* analyzed by RNA-seq. The x-axis indicates fold change in expression and the y-axis indicates the magnitude of expression. The size of dots is positively correlated with the significance of the p-value. Two independent replicates were analyzed with RNA-seq. (**C**) Boxplots showing the expression levels of normal, up-, and down-regulated genes analyzed by reads intensity of RNA-seq of WT and *trbf2*. The p-value was calculated by the t-tests. (**D**) *MA* plot showing the changes of H3K27me3 reads intensity between WT and *trbf2* analyzed by cut&tag. The x-axis indicates the enrichment intensity of each peak and the y-axis indicates log2(difference in H3K27me3 intensity). (**E**) Metagene plots showing H3K27me3 intensity at all H3K27me3 enriched regions (the H3K27me3 enriched regions in WT), the regions gained or lost H3K27me3 in *trbf2* compared with WT. The y-axis indicates reads of exon model per million mapped reads (CPM, read coverage normalized to 1× sequencing depth). (**F**) Heatmaps of H3K27me3 cut&tag reads density at regions that either gained or lost H3K27me3 in *trbf2* compared with WT. Peaks aligned at the peak summit of H3K27me3 are plotted with 5 kb up- and downstream regions (E and F). Two independent replicates were analyzed by cut&tag in (D-F). (**G**) Venn diagram showing the overlaps of DEGs with altered H3K27me3 genes in *trbf2*. The p-value was calculated with hypergeometric tests.

*trbf2* displayed similar developmental defects such as short plant and reduced panicle length as PRC2 mutants including *clf* (Fig. 4A) (*16, 18–20*). We would like to explore the molecular connections between TRBF2 and PRC2. First, we compared the transcriptome profiles of *trbf2* and *clf*. We found 1,479 DEGs in *clf*, including 948 and 531 up- or down-regulated genes respectively (Fig. S9A and table S4). Similar to *trbf2*, most of the up-DEGs in *clf* displayed a repressed expression state in WT (Fig. S9B), consistent with the repression role of PRC2 complex. A total of 375 common DEGs were found in *trbf2* and *clf*, and most of the DEGs displayed the same directions of differential expression (331 out of 375). There were 212 common up-DEGs and 119 common down-DEGs respectively (Fig. S9, C and D). Moreover, changes in expression levels of these genes were positively correlated (Fig. 4B). Thus TRBF2 and PRC2 co-regulate a set of genes. Next, we compared the H3K27me3 levels in *trbf2* and *clf*. Consistent with PRC2 function, most of the altered H3K27me3 regions in *clf* displayed lost H3K27me3 (total, 16,535; lost, 4,479; gained 574) (Fig. S10 and table S5). Further analysis found that only 87 up-regulated regions were common in *trbf2* and *clf* (Fig. 4C), but more than half of the lost H3K27me3 regions in *trbf2* (1,163/2,286) were also decreased in *clf* (Fig. 4C). These data suggest that TRBF2 and PRC2 have commonly targeted chromatin regions and both are required for H3K27me3 deposition at these loci.

**Fig. 4.**
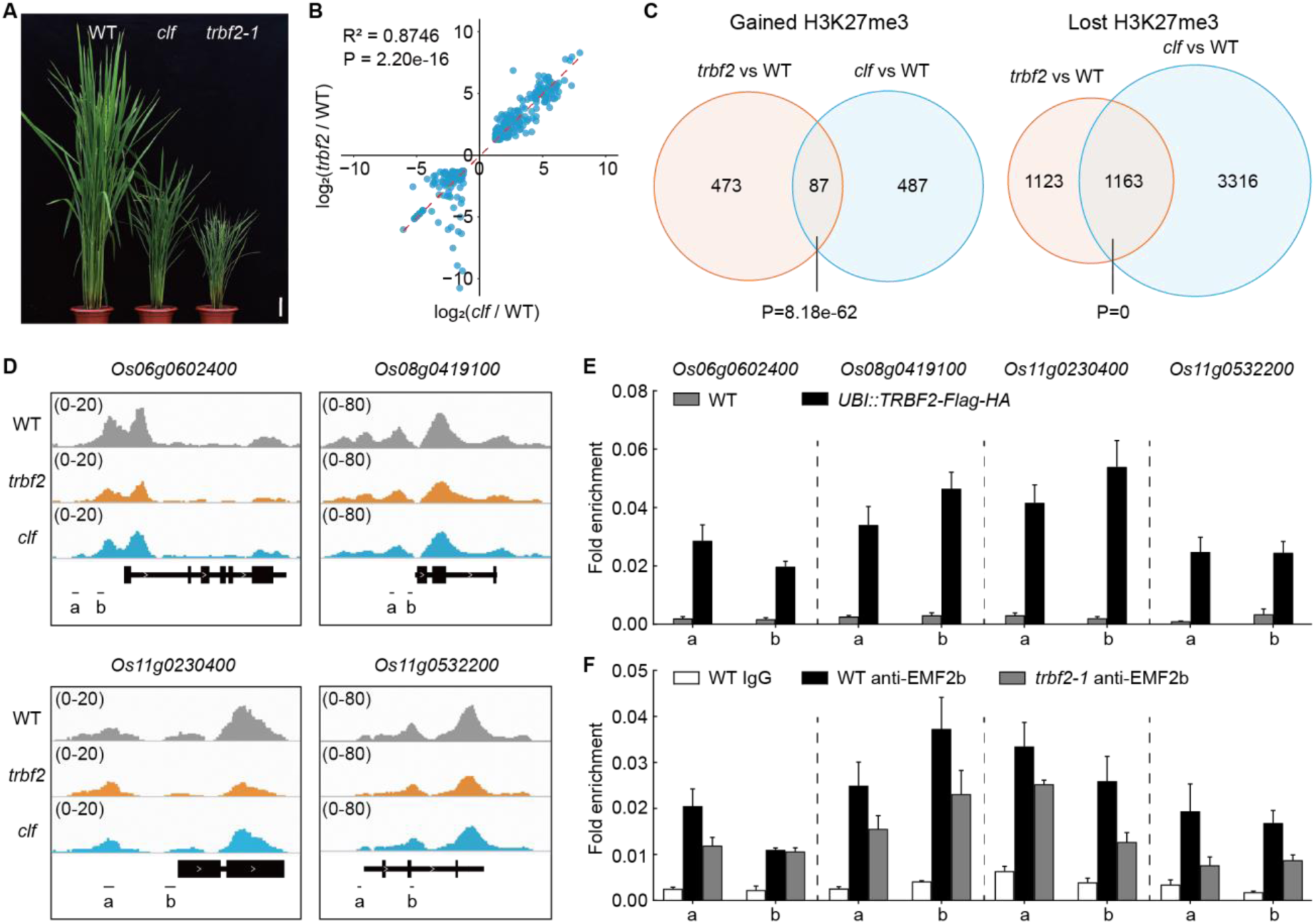
TRBF2 and PRC2 co-regulate gene expression and H3K27me3 modification at target loci. (**A**) Phenotypes of WT, *clf*, and *trbf2* at heading stage. Scale bar, 10 cm. (**B**) Scatter plots showing a positive correlation of up- and down-regulated genes between *trbf2* and *clf*. The line of best fit is shown by the dashed red line, with an adjusted R-value indicated. The p-value was calculated by linear regression. (**C**) Venn diagrams showing the overlaps of gained-or lost-H3K27me3 regions between *trbf2* and *clf*. The p-values were calculated by hypergeometric tests. (**D**) Integrative Genomics Viewer visualization showing lower H3K27me3 levels in *trbf2* and *clf* compared with WT. (**E**) TRBF2 can be enriched at the selected genes measured by anti-Flag ChIP-qPCR. Each examined region, a or b, is shown at the bottom of (D). Values are mean ± SEM (n = 4). (**F**) ChIP-qPCR detection of EMF2b occupancy with anti-EMF2b and IgG (negative control) in WT and *trbf2-1*. Each examined region, a or b, is shown at the bottom of (D). Values are mean ± SEM (n = 3).

Next, we would like to test whether TRBF2 could directly target some of the commonly regulated genes of TRBF2 and CLF. Four genes, which all displayed lower levels of gene expression and H3K27me3 modification both in *trbf2* and *clf*, were selected (Fig. 4D and fig. S11, A to D). The expression levels of these four genes were also de-repressed in the other two *TRBF2* mutants (Fig. S11, E to H). An *UBI::TRBF2-Flag-HA/*WT transgenic line was generated. ChIP-qPCR results showed that TRBF2 could be enriched at the chromatins of these loci (Fig. 4E), indicating these genes are the direct targets of TRBF2. One critical function of phase-separated condensates is to enrich relevant regulators at regulated chromatin regions (*22–25, 29, 30*). So we assayed PRC2 levels at those loci. PRC2 could associate with these loci presented by EMF2b binding levels, and the binding levels were disturbed in *trbf2* (Fig. 4F). Together, our data showed that TRBF2 can concentrate PRC2 at target loci to facilitate H3K27me3 deposition in Polycomb silencing.

In summary, we demonstrate a Polycomb regulation mechanism that involves LLPS in plants. The rice transcription factor TRBF2 forms phase-separated condensates and attracts PRC2 to generate TRBF2-PRC2 aggregates through physical protein interaction. And this mechanism is also conserved in *Arabidopsis* (*13, 26*). TRB proteins are conserved in many plant species, suggesting concentrating PRC2 by TRB-PRC2 aggregates is potential to be widely used in plants. Hence, PRC2 could be concentrated at target loci in gene repression. PRC2 condensates provide an ideal structural hub for chromatin compaction and supply enough space to compact co-regulated chromatin loci together. On the other hand, dense chromatin stimulates PRC2 activity (*31*). Thereby, PRC2 condensation, chromatin compaction, and H3K27me3 three together may generate enough internal force to sustain Polycomb epigenetic memory (*32*). It will now be important to investigate whether the phase-separated TRBF2-PRC2 aggregates contribute to Polycomb body formation and chromatin organization. Transcription factor guided PRC2 targeting mechanism is conserved across plants and animals, but TRB transcription factors are plant-specific. So it is interesting to test whether there is any functional similar transcription factor in non-plant systems.

## Acknowledgments

We thank Professor Liang Chen (Wuhan University) and Professor Shengbo He (South China Agricultural University) for critical comments, Mingliang Tang for microscope support, and members of the Yang research group for helpful discussions. The EMF2b anti-body is a kind gift from Professor Daoxiu Zhou’s Lab (Huazhong Agricultural University). This project was supported by the National Natural Science Foundation of China (31871301), Hubei Hongshan Laboratory, and a start-up fund from Wuhan University.

## Author contributions

H.Y. and H.X. designed the study; H.X. performed and supervised all experimental works; Y, Liu carried out the bioinformatics analysis with help from Y, Li; J.Z. assisted with Co-IP and yeast two-hybrid experiments; N.S. helped to generate plant materials; L.P. and S.L. helped for phenotype measurements; G.X. and Y.Z. helped for phase separation assays; H.Y., H.X. and Y, Liu wrote the manuscript.

## Data and materials availability

The RNA-seq and cut&tag datasets supporting the conclusions of this paper have been deposited in the Sequence Read Archive (SRA) under accession PRJNA811378.

